# Effect of selection on bias and accuracy in genomic prediction of breeding values

**DOI:** 10.1101/298042

**Authors:** G. R. Gowane, Sang Hong Lee, Sam Clark, Nasir Moghaddar, Hawlader A Al-Mamun, Julius H. J. van der Werf

## Abstract

Reference populations for genomic selection (GS) usually involve highly selected individuals, which may result in biased prediction of estimated genomic breeding values (GEBV). In the present study, bias and accuracy of GEBV were explored for various genetic models and prediction methods when using selected individuals for a reference. Data were simulated for an animal breeding program to compare Best Linear Unbiased Prediction of breeding values using pedigree based relationships (PBLUP), genomic relationships for genotyped animals only (GBLUP) and a Single Step approach (SSGBLUP), where information on genotyped individuals was used to infer a matrix **H** with relationships among all available genotyped and non-genotyped individuals that were linked through pedigree. In SSGBLUP, various weights (α=0.95, 0.80, 0.50) for the genomic relationship matrix (**G**) relative to the numerator relationship matrix (**A**) were applied to construct **H** and in another version (SSGBLUP_F), inbreeding was accounted for while computing **A**^-1^. With GBLUP, accuracy of GEBV prediction increased linearly with an increase in the number of animals selected in reference. For the scenario with no-selection and random mating (RR) prediction was unbiased. For GBLUP, lower accuracy and bias observed in the scenarios with selection and random mating (SR) or selection and positive assortative mating (SA), in which prediction bias increased when a smaller and highly selected proportion genotyped. Bias disappeared when all individuals were genotyped. SSGBLUP_F showed higher accuracy compared to GBLUP and bias of prediction was negligible even with selective genotyping. However, PBLUP and SSGBLUP showed bias in SA owing to not fully accounting for allele frequency changes because of selection of quantitative trait loci (QTL) with larger effects and also due to high inbreeding rate. In genetic models with fewer QTL but each with larger effect, predictions were less accurate and more biased for selection scenarios. Results suggest that prediction accuracy and bias is affected by the genetic architecture of the trait. Selective genotyping lead to significant bias in GEBV prediction. SSGBLUP with appropriate scaling of **A** and **G** matrices can provide accurate and less biased prediction but scaling requires careful consideration in populations under selection and with high levels of inbreeding.

## Introduction

Best Linear Unbiased Prediction (BLUP) provides unbiased estimates of breeding values in populations under selection, conditional on the inclusion of all information used in the selection decisions (Henderson 1975; Sorensen and Kennedy 1984). In the case of genomic prediction based on a selected reference population, this condition might not be met, in particular when only genotyped individuals are evaluated in genomic BLUP (GBLUP). A number of studies (VanRaden et al., 2009a, 2009b, Patry and Ducrocq 2011, Vitezica et al. 2011) reported decreased accuracy of genomic estimated breeding value (GEBV) and increased bias due to selective genotyping of sires.

Single Step genomic BLUP (SSGBLUP) (Legarra et al. 2009; Christensen and Lund 2010) combines genomic relationships from genotyped individuals with pedigree relationships with non-genotyped individuals and this integration should allow information on unselected animals to be included, with all relationships tracing back to a conceptual unselected base population. However, SSGBLUP requires the genomic relationship matrix (**G**) and pedigree-based relationship matrix (**A**) to refer to the same base population as otherwise new bias could be introduced. Some studies have discussed this issue and proposed a slight modification in the SSGBLUP procedure (Forni et al. 2011, Vitezica et al. 2011, Christensen et al. 2012). The modification suggested by Vitezica et al. (2011) involved adding the difference between means of **A_22_** and **G** to all elements of **G**, such that the relationships of genotyped individuals are rightly scaled in relation to the base population of all animals in the pedigree. In their study, Vitezica et al. (2011) used a model with only 250 quantitative trait loci (QTL) affecting the trait and 10 generations of assortative mating of sires and dam was used.

The present study aimed at exploring the effect of selection on genomic prediction for a wider range of scenarios. Genetic evaluations based on pedigree BLUP (PBLUP), GBLUP and SSGBLUP were compared. We investigated a number of factors affecting bias and accuracy of genetic evaluations, including (1) the proportion of individuals selected to have genotype information; (2) the genetic structure of the trait as determined by the heritability and the number of QTLs explaining the variance in the trait; and (3) scenarios with and without selection and assortative mating.

## Material and Methods

### Population and genotype simulation

Data were simulated using QMSim (Sargolzaei & Schenkel, 2009). A historical population with effective population size (Ne) of 100 was generated with 50 males and 50 females producing 2 progeny each by random union of gametes in each of 95 generations and thereafter the number of progeny was gradually expanded to 1000 offspring until the 100^th^ generation. In the subsequent 10 generations (101-110), 50 males were mated to 500 females who produced 1000 progeny. The genomic structure consisted of 30 chromosomes of equal length (1 Morgan). Biallelic markers (60,000) were randomly distributed across the genome with an equal frequency (0.5) in the first generation of the historical population. The mutation rate of the markers and QTL was 2.5×10^−8^ per locus per generation (Hickey and Gorjanc 2012). In order to make sure that data were correctly simulated for the historical population, we confirmed whether population parameters like effective number of chromosome segments (M_e_) in the simulated data agreed with the expectation from theory given the value of Ne and the family structure in the population (Lee et al. 2017). M_e_ was determined from the variation in genomic relationship among members of the population. Four genetic models were simulated that differed in the number and distribution of QTL effects; with 90, 990, 9,990 and 60,000 QTLs. The QTL allele effects were sampled from a Gamma distribution with a shape and scale parameter of 0.4 and 1.0, respectively. Genotype effects at individual QTL were aggregated to form true breeding values (TBV) and these were re-scaled to match the input value for the additive genetic variance. For individuals in the 101^st^ generation these breeding values were normally distributed with mean 0 and variance h^2^. Residual effects on phenotype were independent and normally distributed with mean 0 and variance (1-h^2^). Therefore, the mean and variance for the simulated phenotypes were zero and one, respectively. Phenotypes were created for all individuals in the last 10 generations. Three different values for heritability (h^2^ = 0.1, 0.3 and 0.5) were tested for each scenario. Selection of sires was either random or based on estimated breeding values (EBV) obtained by PBLUP. Positive assortative mating was also applied, giving rise to three different scenarios: I: Random selection and random mating (RR), II: Selection on EBV and Random mating (SR), III: Selection on EBV and assortative mating based on EBV (SA). Selection was with non-overlapping generations and in each generation 10% of males were mated to all young females. Under each of these 3 scenarios, predictions of EBV were obtained by each of the different methods and modelling parameters. Scenario PBLUP involved BLUP prediction based on all phenotype and pedigree information from 9 generations to predict EBV in the 10^th^ generation. For each of the last 4 generations (6^th^ to 9^th^ generation), either 125 (25%), 250 (50%) or 500 (100%) males were selected for genotyping to form a reference population consisting of 500, 1000 or 2000 animals (scenario G500, G1000 or G2000). In scenario G9550, all the 9550 animals from all 9 preceding generations were genotyped and used as a reference population. In scenario SS500, SS1000 and SS2000, SSGBLUP was used for prediction of breeding values combining information from pedigree (9550 individual’s pedigree) and genomic relationships from either 500, 1000 or 2000 genotyped individuals in the reference, respectively. For each scenario, we conducted 25 replicates.

### Analysis and estimation of breeding value

In total 1000 selection candidates in the 10^th^ generation were used as a validation population to determine the bias and accuracy of estimated breeding values.

PBLUP analysis was based on pedigree information from the 9 preceding generations that include 9550 animals. Alternatively, PBLUP accounting for inbreeding accumulated since generation 0 while constructing **A** was also used for prediction of breeding values and was named PBLUP_F.

The following linear mixed model was used.

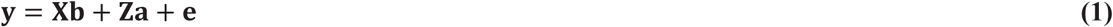

where **y** is a vector of observations; **b** is a vector of fixed effects (sex); **a** is a vector of direct additive genetic effects of individual animals; **e** is a vector of residual errors; and **X** and **Z**, are known incidence matrices. Assumptions in the model were **a** ~ N(0, **A**σ^2^_a_) and **e** ~ N(0, **I**σ^2^_e_); where **A** is the numerator relationship matrix between animals derived from pedigree, **I** is an identity matrix, and σ^2^_a_ and σ^2^_e_ are additive genetic and residual variances, respectively.

GBLUP analysis was used to estimate genomic breeding values. A genomic relationship matrix (**G)** was constructed by method of Yang et al. (2010) using PLINK 1.9 (Chang et al., 2015). The genomic analysis was done using MTG2 (Lee and van der Werf, 2016) to predict GEBVs for animals in generation 10. The model used for analysis was

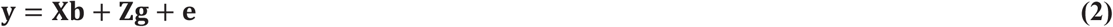

where **g** is a vector of additive genetic effects of the individual animal, with **g** ~ N(0, **G**σ^2^_g_) and other terms are defined as above.

<u>SSGBLUP analysis</u> combined pedigree and genomic information in a single step extending the animal model to include marker genotypes. The model for evaluation was:

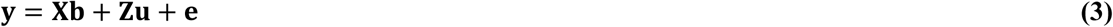

where, **u** is the vector of direct additive genetic effects of individual animals, assumed to be normally distributed with **u** ~ N**(0, Hσ^2^_u_).** The **H** matrix includes both non-genotyped and genotyped individuals (Christensen and Lund, 2010; Aguilar et al. 2010). Other terms are already defined as above. This method did not include inbreeding in the set up of **A**^-1^, however, it considered inbreeding in **A**_22_^-1^. The **H^-1^** derivation was as under

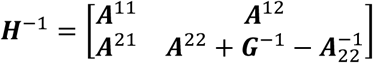

where **A**_22_ is a pedigree-based numerator relationship matrix for genotyped animals (Aguilar et al., 2010).

The tuning of the **H^-1^** matrix component relating to genotyped individuals was according to **A^22^ + (G^*-1^-A_22_^-1^)** with **G^*^** being α**G +** (1-α)**A_22_** and three values of **α** were compared; 0.95, 0.80 and 0.50. In addition to this, the **H** matrix was modified as suggested by Vitezica et al. (2011) that involved adding a small constant to all elements of the **G**. The value was derived as the difference between the means for **A_22_** and **G**. Another single step method (SSGBLUP_F) was employed where **A**^-1^ was correctly constructed taking inbreeding into account. The family of BLUPF90 programs (Misztal, 2008; Aguillar et al. 2014) were used to analyse the data.

Accuracy of prediction of breeding values was obtained as the Pearson correlation between TBV and GEBV of all individuals in generation 10. Bias was estimated as the deviation of regression coefficient of TBV on GEBV from unity.

## Results

### Validation of the simulated data

We showed that in the simulation the observed value for the effective number of chromosome segments (M_e_) agreed with the expected M_e_, given Ne, as described by Lee et al. (2017). When using 30 chromosomes, each with 1 Morgan long, the observed and expected M_e_ were similar and the effects of the number of historical generations and mutation rate were low (Table S1). This indicates that the simulated populations should have properties that align with the pre-defined population parameter for N_e_, which are relatively robust towards the number of historical generations and the assumed mutation rate in the simulation. Furthermore, we confirmed that observed and expected prediction accuracy agreed well (Table S1), validating the reported values for prediction accuracy in our simulation.

### Increasing number of individuals in reference population increases accuracy of GEBV prediction

A larger proportion of genotyped animals led to a larger sample size in the reference population which increased the accuracy of GBLUP as well as for SSGBLUP prediction of GEBV (Table 1). The increase in the accuracy was approximately linear with sample size across the different scenarios where the number of genotyped males increased from 25% to 100% from the last 4 generations.

**Table 1.**
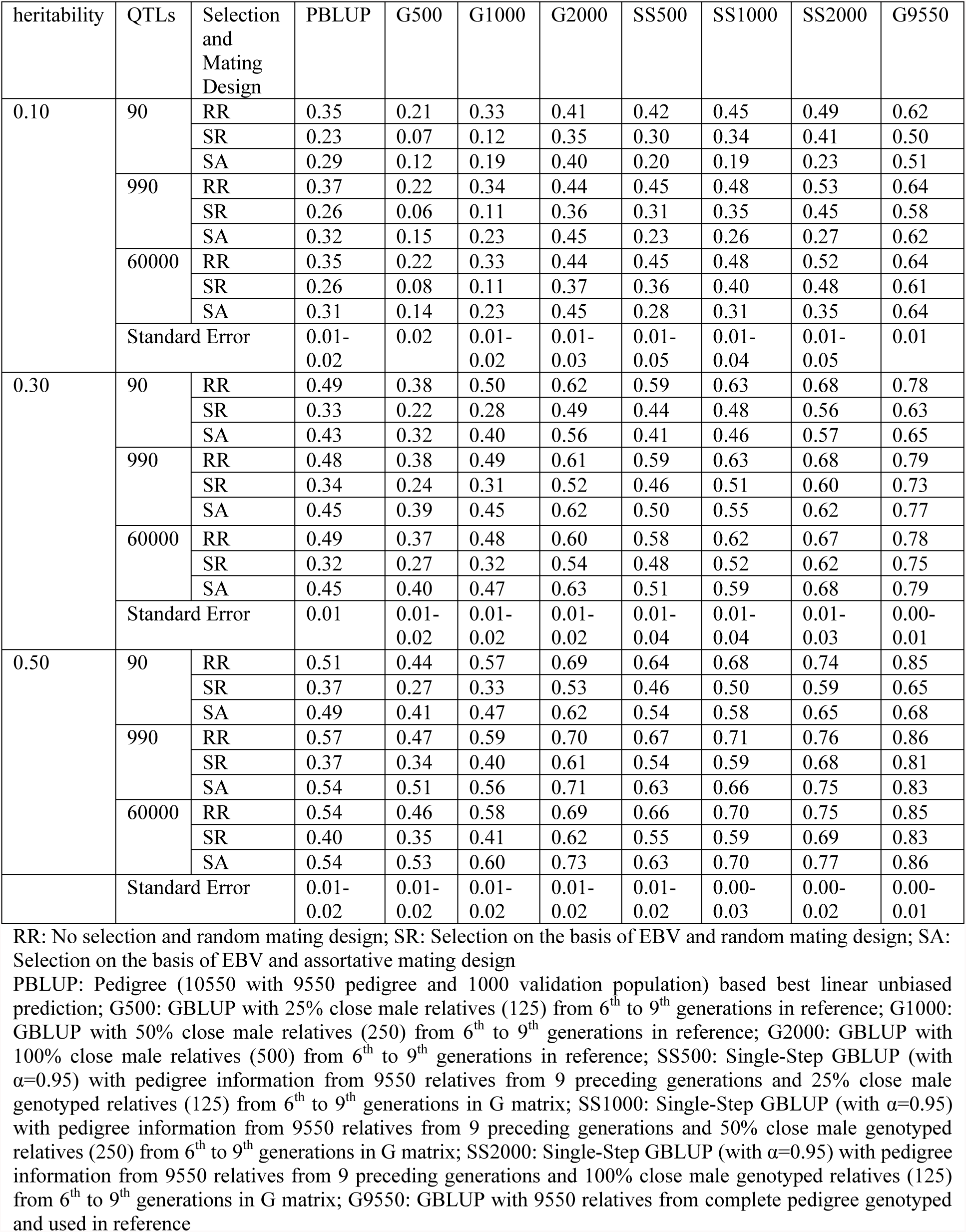
Estimated accuracy of (G)EBV prediction using different methods and scenarios

For the RR scenario and a trait with heritability of 0.3 controlled by 990 QTLs, PBLUP gave an accuracy of 0.48±0.01. The accuracy of G1000 from GBLUP was similar to that of PBLUP. The accuracy increased with a higher sample size *e.g*. 0.61 for G2000 (Table 1). Accuracy with SSGBLUP was much higher; *i.e.* 0.59±0.01 with SS500, increasing to 0.68±0.01 for SS2000. Using genotype information on all animals (G9550) gave the highest prediction accuracy, as expected (0.79±0.01).

### Effect of selection on prediction accuracy

In the SR scenario, the accuracy of both EBV and GEBV were lower than in RR. For PBLUP, the accuracy was 0.34±0.01. A similar value for the prediction accuracy was obtained by GBLUP with G1000 (0.31±0.01). SSGBLUP gave an accuracy of 0.46±0.01 and 0.60±0.01, with SS500 and SS2000, respectively. For the SA scenario, the prediction accuracy from PBLUP was 0.45±0.01. A similar prediction accuracy was obtained with G1000 (0.45±0.02) and a much higher accuracy was obtained by SS1000 (0.55±0.02).

Variation in α for tuning the contribution from the **G**-matrix to form **H** affected the accuracy of prediction in SSGBLUP for all selection and mating designs. Three values of α were tested; 0.95, 0.80 and 0.50. A genetic model with h^2^=0.3, and 4 QTL models (90, 990, 9990, 60000) were compared. For the RR and SR scenarios, using α=0.95 gave the highest accuracy. In the RR scenario, the accuracy decreased by nearly 3.0 to 4.5% and for the SR scenario by 5.0 to 8.0% if α was changed from 0.95 to 0.50. However, for the SA scenario, using α=0.50 gave the highest accuracy. For the 990-QTL scenario with SA, the SS500 model had the accuracy improved from 0.50 ± 0.03 for α =0.95, to 0.51±0.03 for α =0.80 and 0.51 ± 0.03 for α = 0.50 (Table 3). Improvements from changing α from 0.95 to 0.50 were 1.8% for SS1000 and 3.2% for SS2000. For the 90-QTL model, much higher gain in accuracy was observed for the SA scenario by changing α from 0.95 to 0.5 for SS500 (9.8%) and SS1000 (8.7%). For model with a larger number of QTL, the reduction of α did not benefit the accuracy.

**Table 3.** Estimated accuracy and bias of GEBV prediction in Single Step approach with tuning of H matrix

| Criteria | QTLs | Selection and Mating Design | SS500<br>$\alpha=0.95$,<br>\$ | SS1000<br>$\alpha=0.95$,<br>\$ | SS2000<br>$\alpha=0.95$,<br>\$ | SS500<br>$\alpha=0.80$ | SS1000<br>$\alpha=0.80$ | SS2000<br>$\alpha=0.80$ | SS500<br>$\alpha=0.50$ | SS1000<br>$\alpha=0.50$ | SS2000<br>$\alpha=0.50$ | SS500<br>F | SS1000<br>F | SS2000<br>F |
| --- | --- | --- | --- | --- | --- | --- | --- | --- | --- | --- | --- | --- | --- | --- |
| Accuracy | 90 | RR | 0.59 | 0.63 | 0.68 | 0.59 | 0.63 | 0.68 | 0.58 | 0.61 | 0.66 | 0.59 | 0.63 | 0.68 |
|  |  | SR | 0.44 | 0.48 | 0.56 | 0.44 | 0.48 | 0.56 | 0.42 | 0.45 | 0.55 | 0.44 | 0.48 | 0.57 |
|  |  | SA | 0.41 | 0.46 | 0.57 | 0.43 | 0.49 | 0.57 | 0.45 | 0.50 | 0.58 | 0.51 | 0.55 | 0.61 |
|  | 990 | RR | 0.59 | 0.63 | 0.68 | 0.59 | 0.63 | 0.68 | 0.57 | 0.61 | 0.65 | 0.59 | 0.63 | 0.68 |
|  |  | SR | 0.46 | 0.51 | 0.60 | 0.46 | 0.50 | 0.60 | 0.43 | 0.47 | 0.57 | 0.46 | 0.51 | 0.60 |
|  |  | SA | 0.50 | 0.55 | 0.62 | 0.51 | 0.55 | 0.64 | 0.51 | 0.56 | 0.64 | 0.57 | 0.61 | 0.60 |
|  | 60000 | RR | 0.58 | 0.62 | 0.67 | 0.58 | 0.62 | 0.66 | 0.57 | 0.60 | 0.64 | 0.58 | 0.62 | 0.67 |
|  |  | SR | 0.48 | 0.52 | 0.62 | 0.47 | 0.51 | 0.61 | 0.44 | 0.48 | 0.59 | 0.48 | 0.52 | 0.62 |
|  |  | SA | 0.51 | 0.59 | 0.68 | 0.53 | 0.59 | 0.68 | 0.57 | 0.60 | 0.67 | 0.60 | 0.63 | 0.62 |
|  | Standard Error |  | 0.01-0.04 | 0.01-0.04 | 0.01-0.03 | 0.01-0.02 | 0.01-0.02 | 0.01-0.02 | 0.01-0.03 | 0.01 | 0.01-0.02 | 0.01-0.02 | 0.01-0.02 | 0.01-0.02 |
| Bias | 90 | RR | 1.00 | 0.99 | 1.00 | 1.04 | 1.03 | 1.04 | 1.10 | 1.11 | 1.13 | 1.03 | 1.01 | 1.02 |
|  |  | SR | 0.85 | 0.86 | 0.80 | 0.90 | 0.92 | 0.87 | 0.96 | 1.00 | 1.00 | 0.90 | 0.91 | 0.86 |
|  |  | SA | 0.49 | 0.53 | 0.56 | 0.54 | 0.58 | 0.60 | 0.65 | 0.69 | 0.70 | 0.81 | 0.82 | 0.81 |
|  | 990 | RR | 1.01 | 1.00 | 1.00 | 1.04 | 1.05 | 1.04 | 1.09 | 1.11 | 1.23 | 1.03 | 1.03 | 1.02 |
|  |  | SR | 1.00 | 1.02 | 0.96 | 1.05 | 1.08 | 1.03 | 1.12 | 1.16 | 1.18 | 1.06 | 1.08 | 1.02 |
|  |  | SA | 0.68 | 0.72 | 0.70 | 0.74 | 0.78 | 0.76 | 0.87 | 0.90 | 0.87 | 1.03 | 1.04 | 1.02 |
|  | 60000 | RR | 0.97 | 0.97 | 0.97 | 1.00 | 1.01 | 1.01 | 1.05 | 1.07 | 1.09 | 0.99 | 1.00 | 0.99 |
|  |  | SR | 1.02 | 1.04 | 0.97 | 1.08 | 1.11 | 1.05 | 1.16 | 1.20 | 1.21 | 1.09 | 1.10 | 1.03 |
|  |  | SA | 0.73 | 0.79 | 0.77 | 0.79 | 0.84 | 0.82 | 0.95 | 0.97 | 0.94 | 1.08 | 1.09 | 1.03 |
|  | Standard Error |  | 0.02-0.06 | 0.01-0.05 | 0.01-0.04 | 0.02-0.04 | 0.01-0.04 | 0.01-0.03 | 0.02-0.04 | 0.02-0.04 | 0.01-0.04 | 0.01-0.03 | 0.01-0.04 | 0.01-0.03 |

The simulated data with $h^2 = 0.3$ were used; \$: modification of SSGBLUP as suggested by Vitezica et al. (2011), $\alpha=0.50$ : shrinking information from **G** matrix to 50%, $\alpha=0.95$ : shrinking information from **G** matrix to 95%, F: SSGBLUP\_F accounting for inbreeding in $\mathbf{A}^{-1}$ and $\mathbf{A}_{22}^{-1}$ .

### Accounting for Inbreeding in **A**^-1^ increases accuracy of Single Step prediction for populations under selection

Accumulation of inbreeding was higher in a selection scenario with assortative mating (Table 5). In the validation population, inbreeding was 2.3% in RR, 3.7% in SR and 9.5% in the SA scenario. For the SA scenario, SSGBLUP (without inbreeding) did not help improve accuracy much over GBLUP. Including inbreeding while constructing **A**^-1^ (SSGBLUP_F) resulted in more accurate estimates. For the SS500 we observed a strong improvement in accuracy if inbreeding was accounted for in the genetic evaluation in single step. For the 990QTL model (h^2^=0.3), accuracy of SS500 increased from 0.50 (SSGBLUP) to 0.57 (SSGBLUP_F) (Table 3).

**Table 5.**
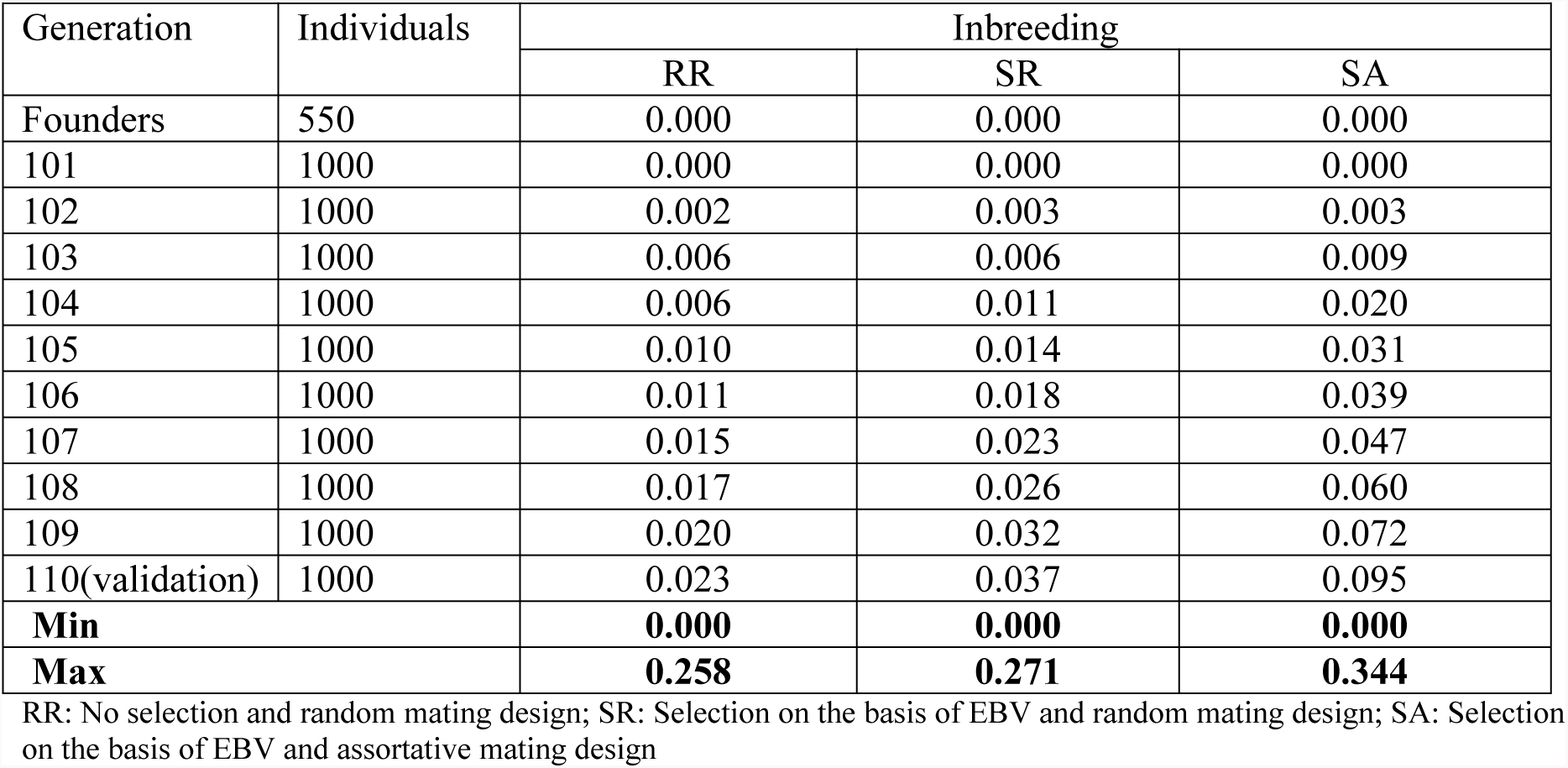
The distribution of inbreeding coefficietns in generations for different selection and mating design scenarios

### Effect of different heritability estimates and QTL models on prediction accuracy

With h^2^ = 0.1, the accuracy of prediction was significantly lower across all scenarios for PBLUP, GBLUP and SSGBLUP. For PBLUP, a change of h^2^ from 0.3 to 0.5 increased the accuracy by 4.1 to 18.8%, 8.8 to 25.0% and 14.0 to 22.7% for RR, SR and SA scenarios, respectively, across the different models. Similarly, a gain in accuracy was observed with increasing h^2^ for GBLUP and SSGBLUP across all different genetic models and scenarios. For GBLUP, the magnitude of increase in accuracy with higher heritability was larger, compared to PBLUP and SSGBLUP and more so when a smaller proportion was genotyped. Also for the scenarios with selection (SR and SA) a higher gain in accuracy compared to the RR scenario was observed with higher heritability models.

Variation in the number of QTLs did not affect the accuracy of prediction of GEBV for the RR scenario. Similarly, accuracy for EBV obtained by PBLUP was unaffected by variation in QTL model for RR, SR and SA scenarios. Prediction accuracy was affected by QTL number for SR and SA scenarios when using GBLUP and SSGBLUP, where the prediction accuracy improved when the number of QTL in the simulated genetic model increased, with more gains for SA than in the SR scenario (Table 1).

### Effect of selective genotyping on the bias of GEBV prediction

In a genetic model with h^2^=0.3 and 990-QTL, all methods showed no bias with regression coefficients around one for no-selection and random mating (RR) scenario (Table 2). Bias of prediction was a common feature in GBLUP with selective genotyping (G500, G1000 and G2000) in SR and SA design with either under-dispersed or over-dispersed regression coefficients (Table 2). The bias reduced when a larger proportion of males was genotyped, which also led to more genotyped individuals in the reference population. For the SR design, bias in GEBV prediction was evident. G500 and G1000 were most biased with regression coefficients of 0.78±0.06 and 0.85±0.03, respectively. GBLUP with G2000 was less biased (0.96±0.02). For the SA design, the GBLUP approach gave slightly under-dispersed GEBVs with the regression coefficient equal to 1.11±0.07 and 1.13±0.05 for G500 and G1000, respectively. However, for G2000 and G9550, the estimated regression coefficient was 1.01±0.03 and 0.96±0.01, respectively, indicating less bias when genomic information is available on more or on all animals.

**Table 2.** Bias of (G)EBV prediction using different methods and scenarios

| heritability | QTLs | Selection and Mating Design | PBLUP | G500 | G1000 | G2000 | SS500 | SS1000 | SS2000 | G9550 |
| --- | --- | --- | --- | --- | --- | --- | --- | --- | --- | --- |
| 0.10 | 90 | RR | 0.99 | 0.65 | 1.09 | 0.99 | 0.93 | 0.93 | 0.93 | 0.99 |
|  |  | SR | 0.88 | 1.49 | 1.45 | 0.93 | 0.81 | 0.82 | 0.77 | 0.82 |
|  |  | SA | 0.76 | 0.18 | 0.51 | 0.83 | 0.26 | 0.25 | 0.24 | 0.77 |
|  | 990 | RR | 1.08 | 0.77 | 1.15 | 1.04 | 1.00 | 1.01 | 1.00 | 1.02 |
|  |  | SR | 1.08 | -0.1 | 0.37 | 0.90 | 0.88 | 0.92 | 0.89 | 0.99 |
|  |  | SA | 0.88 | 0.82 | 1.00 | 0.97 | 0.32 | 0.33 | 0.32 | 0.94 |
|  | 60000 | RR | 1.02 | 2.31 | 1.40 | 1.09 | 1.00 | 1.01 | 0.99 | 1.01 |
|  |  | SR | 1.14 | 0.60 | 0.46 | 0.87 | 1.05 | 1.06 | 0.96 | 1.02 |
|  |  | SA | 0.92 | 0.38 | 0.82 | 0.97 | 0.46 | 0.46 | 0.41 | 0.98 |
|  | Standard Error |  | 0.03-0.05 | 0.10-1.61 | 0.07-2.20 | 0.03-0.13 | 0.03-0.07 | 0.02-0.07 | 0.02-0.06 | 0.01-0.02 |
| 0.30 | 90 | RR | 1.04 | 0.97 | 1.03 | 1.02 | 1.00 | 0.99 | 1.00 | 1.00 |
|  |  | SR | 0.87 | 0.67 | 0.73 | 0.84 | 0.85 | 0.86 | 0.80 | 0.80 |
|  |  | SA | 0.72 | 0.81 | 0.96 | 0.90 | 0.49 | 0.53 | 0.56 | 0.78 |
|  | 990 | RR | 0.95 | 1.16 | 1.01 | 1.01 | 1.01 | 1.00 | 1.00 | 1.00 |
|  |  | SR | 1.00 | 0.78 | 0.85 | 0.96 | 1.00 | 1.02 | 0.96 | 0.98 |
|  |  | SA | 0.88 | 1.11 | 1.13 | 1.01 | 0.68 | 0.72 | 0.70 | 0.96 |
|  | 60000 | RR | 0.99 | 1.02 | 1.00 | 1.00 | 0.97 | 0.97 | 0.97 | 1.01 |
|  |  | SR | 0.93 | 0.84 | 0.87 | 0.96 | 1.02 | 1.04 | 0.97 | 1.00 |
|  |  | SA | 0.91 | 1.26 | 1.17 | 1.04 | 0.73 | 0.79 | 0.77 | 0.99 |
|  | Standard Error |  | 0.01-0.03 | 0.04-0.12 | 0.03-0.06 | 0.01-0.03 | 0.01-0.06 | 0.01-0.05 | 0.01-0.03 | 0.01-0.02 |
| 0.50 | 90 | RR | 0.93 | 0.97 | 0.98 | 0.98 | 0.95 | 0.96 | 0.97 | 0.99 |
|  |  | SR | 0.75 | 0.67 | 0.74 | 0.84 | 0.76 | 0.79 | 0.75 | 0.75 |
|  |  | SA | 0.78 | 0.85 | 0.94 | 0.87 | 0.63 | 0.67 | 0.67 | 0.75 |
|  | 990 | RR | 1.02 | 1.07 | 1.04 | 1.04 | 1.00 | 1.00 | 1.00 | 1.01 |
|  |  | SR | 0.91 | 0.85 | 0.92 | 0.97 | 1.02 | 1.06 | 0.97 | 0.98 |
|  |  | SA | 0.93 | 1.09 | 1.13 | 1.01 | 0.86 | 0.88 | 0.85 | 0.96 |
|  | 60000 | RR | 0.99 | 1.05 | 1.01 | 0.99 | 0.98 | 0.98 | 0.98 | 1.00 |
|  |  | SR | 1.01 | 0.88 | 0.96 | 0.99 | 1.05 | 1.10 | 1.00 | 1.00 |
|  |  | SA | 0.95 | 1.15 | 1.23 | 1.05 | 0.83 | 0.91 | 0.88 | 1.00 |
|  | Standard Error |  | 0.01-0.03 | 0.02-0.05 | 0.02-0.03 | 0.01-0.02 | 0.01-0.05 | 0.01-0.04 | 0.01-0.03 | 0.01-0.02 |

Footnotes for Table 2 are the same as in Table 1.

### Effect of selection and mating designs on bias of GEBV prediction

For the SR scenario, the bias of prediction was removed with the SSGBLUP under the 990-QTL model but some bias was observed under the 90-QTL model. However, bias with SSGBLUP was significantly smaller than with GBLUP, especially when only a highly selected proportion was genotyped. For the SA design, the bias in the prediction of breeding value was very high for all methods of prediction, including PBLUP, where a regression coefficient of 0.88±0.02 was obtained. The SSGBLUP methods using α=0.95 was more biased in the SA scenario. For SS500, regression was 0.68±0.05, for SS1000 it was 0.72±0.05 and even for SS2000 model, the regression was 0.70±0.04 and bias was larger with SSGBLUP than with GBLUP.

Reducing the weight given to the **G-**matrix (α from 0.95 to 0.80 and then to 0.50) in SSGBLUP, increased the regression coefficient for the SA scenario, i.e. there was less bias. The improvement in regression coefficient for the 990-QTL (h^2^=0.3) model by shifting α from 0.95 to 0.50 was 27.9% for SS500, 25% for SS1000 and 24.3% for SS2000 (Table 3). One interesting feature with the SA scenario is that by decreasing the weight of **G** matrix (α from 0.95 to 0.5) in SSGBLUP method also improved the accuracy (Table 3), although the difference was statistically non-significant. The SR scenario showed almost no reduction in accuracy when lowering the value for α.

### Including inbreeding in **A**^-1^ for SSGBLUP reduces bias of prediction of GEBV in population under selection

Bias of prediction for the SA scenario was high, even with the SSGBLUP method. However, the bias was practically removed when accounting for inbreeding in the **A** matrix (SSGBLUP_F) (Table 3). For the RR scenario, all estimates were nearly unbiased and inclusion of inbreeding in **A**^-1^ did not affect the GEBV prediction. For the SR scenario, over-dispersed GEBV (regression coefficients>1) were obtained for SSGBLUP(α=0.50), however, SSGBLUP_F resulted in regression coefficients close to 1, and at par with PBLUP. For the SA scenario, a tremendous improvement in bias of prediction was observed when inbreeding was accounted for in **A**. For the 90QTL scenario, regression coefficient was 0.81 for SS500 (SSGBLUP_F) as compared to 0.69 for SS500(α=0.50), 0.72 for PBLUP and 0.76 for PBLUP including inbreeding (PBLUP_F). This improvement was mainly due to incorporation of inbreeding while constructing **A**^-1^. PBLUP estimations without including inbreeding were similar to PBLUP including inbreeding, although little gain in regression was observed for SA scenario that was in-significant. It is therefore important for the SSGBLUP method to incorporate inbreeding than for the BLUP method, because **G**^-1^ will automatically account for inbreeding, and to be consistent with **A**, **A**^-1^ needs to account for it as well while constructing **H**^-1^.

### Effect of heritability and QTL number variation on bias of prediction

With a higher heritability (increasing from 0.3 to 0.5), more accurate predictions with lower bias were obtained across prediction methods and mating designs. However, with a low heritability (h^2^ = 0.1), the prediction was significantly biased for GBLUP or Single Step methods (Table 2). For the RR scenario, whether using a small or large number of QTLs, there was not much bias observed. However, when selection was involved (SR and SA scenarios), the 90-QTL model resulted in significantly biased predictions for all the methods, including PBLUP and G9550 while models with higher numbers of QTL resulted in lower bias of prediction (Table 2). This happened mainly due to selection of QTL with large effect.

## Discussion

In the pedigree-based genetic evaluation of a breeding program, it is assumed that the individuals in the base population are unselected and unrelated having average inbreeding coefficient of zero (Falconer, 1996). Henderson (1975) showed that under the infinitesimal genetic model all subsequent selection is conditional upon the unselected base population and is accounted for in BLUP prediction, provided all data is included in the evaluation model. When genomic prediction is based on the genotyped animals alone, this condition is not met, and this gives rise to biased predictions as was shown in the GBLUP scenarios in this study. The bias was manifested in an over-dispersion of the GEBVs. In other words, GEBVs are over-predicted for the top animals at the moment of selection. Bias was smaller when selection for genotyping was less intense and bias was removed when all data that was used in selection decisions was included, as was the case in the SSGBLUP method.

Results in our study showed that all prediction methods had lower accuracy in the scenarios with selection (SR and SA). Even the PBLUP method showed a significant (20-30%) decrease in accuracy for all selection scenarios. The lower accuracy in the SR and SA scenarios compared to RR is likely due to the Bulmer effect, *i.e.* the variation between families is reduced due to selection, leading to a reduction of genetic variance and a lower correlation between EBV and TBV (Bijma, 2012). This effect will be relatively large in our study, where the accuracy was validated in the animals from the last generation that had no phenotypic records themselves and the EBV was largely based on information through either pedigree or genomic relationships. The decrease in accuracy was lower for the SSGBLUP compared with the PBLUP scenarios, as the GEBV is more based on information within families (Clark et al. 2013). Assortative mating increases the variance among offspring and shows therefore higher correlations between EBV and TBV. GBLUP prediction accuracy is likely less affected by the Bulmer effect. However, the accuracy in GBLUP was negatively affected by the effect of bias due to only using genotype information on selected animals. With a larger reference population the effect of selective genotyping is smaller, but also leads to more information being used from genotyped individuals leading to more of the within family information being captured and less selection bias.

The lower accuracies found in present study for GBLUP compared to SSGBLUP were also reported by Vitezica et al. (2011) and can be attributed to SSGBLUP also using information on un-genotyped individuals that are linked through the pedigree. Comparing G2000 and SS2000 results, we found that the gain in the accuracy was small, which similarly was observed across different scenarios. This was probably because most of the information was already captured by **G** consisting of the individuals from the last 4 generations. Although more deep pedigree information is used in the SS2000 model, the ancestral coefficient of relationship used is numerically very small, giving not much gain in accuracy. However, omitting relationships to the unselected base population from the analysis can still have a significant effect on the ability to correct for selection bias, as was demonstrated by Van der Werf and De Boer (1990).

In the present study, data was generated from a population consisting of 50 sires mated to 500 dams, each dam producing 2 progeny. Thus individuals have information on 18 half-sibs and one full-sib, resulting in an estimated N_e_-value for the reference population of approximately 358 (Lee et al. 2017). With an N_e_-value of 358, the expected value for prediction accuracy derived from theory (Lee et al. (2017)) approximately agreed with the observed accuracies in this simulation study (Table S2). For example, the expected accuracies for the scenarios with h^2^=0.3, and with 50000 SNP according to Lee et al. (2017) were 0.385 (G500), 0.508 (G1000), 0.641 (G2000) and 0.877 for G9550. The observed accuracy (under the no-selection scenario) as obtained with GBLUP for the 990-QTL model were 0.38±0.01 (G500), 0.49±0.01 (G1000), 0.61±0.01 (G2000) and 0.79±0.01 (G9550). It is noted that the population structure is more complicated than the full- and half-sib relationships within one generation, which may explain the small difference between the observed and expected prediction accuracy.

Accuracy for GEBV obtained by GBLUP was affected by the assumed QTL model. This was probably due to the fact that the gamma distribution employed in the simulation resulted in a few QTL with large effect. In the validation population (Generation 110) the 90-QTL model (h^2^=0.3) had on average 5.12 ± 0.35 QTL explaining more than 5% variance individually for the RR scenario. This number was 2.68 ± 0.28 for SR and 1.6 ± 0.26 for the SA scenario (Table S3). For the 990-QTL model, QTL explaining more than 5% variance were 0.36 ± 0.11 for RR; 0.2 ± 0.08 for SR and 0.16 ± 0.07 for the SA scenario. Two important things are inferred from above data. First, the number of large QTL is larger in the 90-QTL model as compared to the models with ≥990-QTL, mainly due to sharing of TBV over a large number of QTL in the latter models. Second, there is a loss of QTL with large effects in selection scenarios as compared to RR, thus reducing accuracy in selection scenarios. Selection scenarios reduced the genetic as well as phenotypic variance in validation population as compared to first generation (Table 6), however, it was seen that the reduction in variance was significantly higher for the 90-QTL model compared to higher QTL-models. For h^2^=0.3, the loss of genetic variance was 40% for SR and 60% for the SA scenario in 90-QTL model, whereas it was 24% and 36.1% for the 990-QTL model and the reduction was 16.7% and 28.6% for the 60000-QTL model. Loss of a few large loci due to selection might therefore affect the accuracy as well as bias in 90-QTL model significantly.

**Table 6.** Estimates of phenotypic and genetic variance in the first generation and validation population of the 10^th^ generation

| Scenario | *QTL model | h<sup>2</sup>=0.1 |  |  |  | h<sup>2</sup>=0.3 |  |  |  | h<sup>2</sup>=0.5 |  |  |  |
| --- | --- | --- | --- | --- | --- | --- | --- | --- | --- | --- | --- | --- | --- |
|  |  | Phenotypic variance |  | Genetic variance |  | Phenotypic variance |  | Genetic variance |  | Phenotypic variance |  | Genetic variance |  |
|  |  | G1<br>(0.03-0.05) | G10<br>(0.03-0.05) | G1<br>(0.01) | G10<br>(0.01) | G1<br>(0.03-0.06) | G10<br>(0.04-0.06) | G1<br>(0.01-0.03) | G10<br>(0.02-0.04) | G1<br>(0.04-0.07) | G10<br>(0.04-0.08)) | G1<br>(0.03-0.06) | G10<br>(0.02-0.09) |
| RR | 90 | 1.02 | 0.99 | 0.10 | 0.10 | 0.99 | 0.99 | 0.30 | 0.30 | 1.00 | 0.95 | 0.50 | 0.47 |
|  | 990 | 1.00 | 1.00 | 0.10 | 0.10 | 1.01 | 1.00 | 0.30 | 0.30 | 0.98 | 0.99 | 0.49 | 0.50 |
|  | 60000 | 1.00 | 0.99 | 0.10 | 0.10 | 0.99 | 0.99 | 0.30 | 0.30 | 1.00 | 0.98 | 0.49 | 0.48 |
| SR | 90 | 1.00 | 0.98 | 0.10 | 0.07 | 1.00 | 0.86 | 0.30 | 0.18 | 0.99 | 0.72 | 0.49 | 0.22 |
|  | 990 | 1.02 | 0.98 | 0.10 | 0.08 | 1.00 | 0.94 | 0.29 | 0.22 | 0.99 | 0.87 | 0.50 | 0.38 |
|  | 60000 | 1.01 | 1.00 | 0.10 | 0.09 | 1.00 | 0.95 | 0.30 | 0.25 | 1.00 | 0.90 | 0.50 | 0.39 |
| SA | 90 | 1.00 | 0.95 | 0.10 | 0.06 | 1.03 | 0.85 | 0.35 | 0.14 | 1.14 | 0.70 | 0.64 | 0.20 |
|  | 990 | 1.00 | 0.98 | 0.10 | 0.08 | 1.05 | 0.92 | 0.36 | 0.23 | 1.13 | 0.88 | 0.62 | 0.38 |
|  | 60000 | 1.01 | 0.98 | 0.10 | 0.08 | 1.03 | 0.94 | 0.35 | 0.25 | 1.14 | 0.93 | 0.63 | 0.39 |

Numbers in the parentheses are estimates for range of Standard Deviation (SD)
RR: No selection and random mating design; SR: Selection on the basis of EBV and random mating design; SA: Selection on the basis of EBV and assortative mating design
\*QTL model: number of QTLs explaining the variance of the trait
G1: Generation 1 (First) with 1000 individuals, G10: Validation population in the 10<sup>th</sup> Generation (N=1000)

Accuracy was also affected by allele frequency changes, which were observed from generation 101 to generation 110 mostly in selection scenarios (SR and SA). Allele frequency changes were very high for the 90-QTL model. For the 990-QTL model, allele frequency changed but to a lesser extent. Allele frequencies of the five largest QTL (Table S4) revealed that for the 90-QTL model the change of allele frequency in the selection scenario was very high for QTL with large effect. Thus, in SSGBLUP, it negatively affected accuracy in smaller reference, as **A** does not accommodate allele frequency changes.

Genomic selection exploits more of the Mendelian sampling variance because it used realized rather than expected relationships (Goddard and Hayes, 2007; Clark et al, 2013). Figure 1 shows that genomic relationships have a normal distribution and there are many negative values. The expected relationships based on pedigree have a skewed distribution with only positive values. Table 4 shows that nearly 50% of the elements in **A** with zero or near zero values are actually negative relationships in the **G** matrix. This helps to explain why pedigree based prediction is less accurate than genomic based predictions. The G9550 have genomic information for all the pedigree that also include the base allelic frequency. This may be the reason that the G9550 scenario gave the most accurate and unbiased prediction of GEBV. Sorensen and Kennedy (1984) and Kennedy et al., (1988), emphasized the fact that the covariances of TBVs for selected individuals are not described well enough by **A** (or **G** for instance with GBLUP), unless all records used in selection are accounted for, as happens for pedigree based estimations. As most of the genomic selection programmes use a **G**-matrix that is obtained from individuals from recent generations, the GEBV estimates are usually biased mostly due to omitting information on selection.

**Table 4.** The distribution of pair-wise relationships for the individuals from the GRM and NRM in different selection and mating designs

| #Genetic relationship | RR |  | SR |  | SA |  |
| --- | --- | --- | --- | --- | --- | --- |
|  | GRM | NRM | GRM | NRM | GRM | NRM |
| -0.25 to 0.00 | 302,20,919 | 141,09,673 | 303,08,050 | 144,79,933 | 305,74,102 | 147,68,775 |
| 0.001 to 0.25 | 253,25,866 | 414,06,656 | 252,37,460 | 410,32,352 | 249,62,864 | 403,93,558 |
| 0.251 to 0.50 | 87,462 | 1,13,318 | 88,798 | 1,16,840 | 96,957 | 4,59,724 |
| 0.501 to 0.75 | 11,711 | 16,313 | 11,656 | 16,845 | 12,006 | 23,520 |
| 0.751 to 0.85 | 17 | 15 | 11 | 5 | 44 | 335 |
| 0.851 to 0.95 | 0 | 0 | 0 | 0 | 2 | 58 |
| >0.95 (up to 0.97) | 0 | 0 | 0 | 0 | 0 | 5 |

The total number of pair-wise relationships was 55645975
RR: No selection and random mating design; SR: Selection on the basis of EBV and random mating design; SA: Selection on the basis of EBV and assortative mating design
GRM: genomic relationship matrix, NRM: numerator relationship matrix

Using only selected animals for a reference population can cause biased estimates in GBLUP. The Single Step approach using a **H**-matrix that combines information on **A** and **G** allows to obtain more accurate and less biased estimates of GEBV. The method generally removed the bias seen in GBLUP, although this was not always the case for the SA scenario. Incompatibility of **A** and **G** due to different bases is a thing of concern. In the selection scenario, where highly selected parents or relatives are chosen for constructing reference, the base population frequencies are usually not traceable. We observed in SSGBLUP that by keeping α=0.95 and by using the methods of Vitezica et al. (2011), the bias still existed for the SA scenario. Vitezica et al. (2011) proposed a modification for tuning of **H** matrix for populations under selection that involved fitting a constant to all elements of the **G** matrix that they derived by equating the sum of the elements of the **G** to the sum of the elements of the **A**. Hsu et al. (2017) showed that under selection, if genotypes (SNPs) include QTLs, accuracy and bias of genomic prediction is compromised for Single Step unless the mean of unselected individuals is fitted in the model as a fixed effect. However if the observed SNPs are only markers, the accuracy of prediction may not be improved by the modification proposed by Vitezica et al. (2011). In our data none of the markers (SNPs) were also QTL, although they were generally in LD with QTL.

For the large group of non-genotyped animals, breeding values are, a priori, conditioned on genetic values of genotyped animals (Legarra et al. 2009) that are actually based on current genotypic frequencies of recent generations of selected animals, where significant changes in allelic frequency took place due to selection and assortative mating design. It seems likely that the bias with the SSGBLUP methods is caused by an inconsistent scaling of the **A** and **G**, due to changing allele frequencies with selection. The accumulation of inbreeding in the populations under selection has an effect on the relationship structure in the population. Not all large scale pedigree based genetic evaluation programmes account for inbreeding as it has non-significant influence on the estimations (Meharabani-Yeganeh et al. 2000). However, not accounting for inbreeding when deriving relationship will have an effect on the scaling of **A** versus **G,** genomic relationships automatically account for inbreeding, and this lack of correct scaling can lead to bias of prediction. An inappropriate scaling of **G** versus **A** is also evident from the decreased bias that was observed when the α-value was decreased from 0.95 to 0.5 (Table 3), i.e. when the genomic relationships are given less weight. However, the bias was removed when inbreeding was incorporated while constructing **A**^-1^. Poor performance of SSGBLUP compared to SSGBLUP_F was therefore due to the presence of highly inbred individuals in data but ignoring inbreeding in the prediction model (Meharabani-Yeganeh et al. 2000, Garcia-Baccino et al. 2017). SSGBLUP is more sensitive to this than PBLUP, which is likely due to the need to combine a relationship matrix **G** that accounts for inbreeding implicitly with a pedigree derived matrix that may not account for it. Similarly, BLUP is relatively robust against deviations from the infinitesimal model (Maki-Tanila and Kennedy 1986), but when **G** and **A** need to be combined, the effect of allele frequency changes seem to be larger. We observed more bias for models with fewer QTL where allele frequency changes are more pronounced.

## Conclusion

This study showed that genomic selection using only highly selected genotyped individuals in the reference for genomic prediction results in considerable bias. The Single Step approach resulted in more accurate and less biased estimates of breeding value because it also takes into account the information from non-selected and non-genotyped individuals. However, with selection and assortative mating, some bias was also observed with the Single Step method, likely due to inappropriate merging of the **A** and **G**-matrices due to allele frequency changes of large QTL as a result of selection and also due to ignoring inbreeding in building **A^-1^**. We conclude therefore that the Single Step approach can easily cause bias as it is quite sensitive to inappropriate scaling of **A** and **G**-matrices, especially with selection and selective genotyping, and with considerable rates of inbreeding, but bias can be minimized when scaling is appropriate.

## Acknowledgements

GRG duly acknowledge the support provided by Indian Council of Agricultural Research (ICAR) and University of New England Australia, and also financial support provided by Endeavour Research Fellowship (Australia). Dr. Rohan L. Fernando (Iowa State University, USA) and Dr. Andres Legarra (INRA, France) are gratefully acknowledged for the useful discussions. SHL is an Australian Research Council Future Fellow (FT160100229).

## Data availability statement

The authors affirm that all data necessary for confirming the conclusions of the article are present within the article, figure, and tables.

---

**Table S1:** Validation for the simulated test data with theory (Lee et al. 2017)

| Test Data X |  |  |  |  |  |  |
| --- | --- | --- | --- | --- | --- | --- |
| Parameters | Scenario1 | Scenario2 | Scenario3 | Scenario4 | Scenario5 | Scenario6 |
| Ne | 100 | 100 | 100 | 500 | 500 | 500 |
| Generations | 50 | 100 | 200 | 250 | 500 | 1000 |
| Observed Me±SE | 246.3±1.54 | 242.2±1.37 | 230.4±1.31 | 1016.8±1.85 | 988.8±2.00 | 940.3±1.72 |
| Expected Me | 246 |  |  | 1157 |  |  |
| Test Data Y |  |  |  |  |  |  |
| Parameters | Scenario1 | Scenario2 | Scenario3 | Scenario4 | Scenario5 | Scenario6 |
| Ne | 100 | 100 | 100 | 500 | 500 | 500 |
| Generations | 100 | 100 | 100 | 500 | 500 | 500 |
| Mutation Rate | 2.5x10<sup>-3</sup> | 2.5x10<sup>-5</sup> | 2.5x10<sup>-8</sup> | 2.5x10<sup>-3</sup> | 2.5x10<sup>-5</sup> | 2.5x10<sup>-8</sup> |
| Observed Me±SE | 272.9±1.38 | 246.2±1.09 | 248.3±0.82 | 1171±2.69 | 1006.5±0.74 | 1007.2±1.53 |
| Expected Me | 246 |  |  |  | 1157 |  |
| Test Data Z |  |  |  |  |  |  |
| Observed Me±SE | Expected Me |  | Observed accuracy |  | Expected accuracy |  |
| 232.36±1.42 | 246 |  | 0.818±0.003 |  | 0.818 |  |

In the test data X, variations included effective population size (Ne) of 100 for historical population with generations 50, 100 and 200, similarly for Ne of 500, historical generations of 250, 500 and 1000 were simulated. 100 replicates of test data X were run to obtain Me. In test data Y, the sensitivity to the mutation rate was analysed for historical populations with three different mutation rates namely high ( $2.5 \times 10^{-3}$ ), medium ( $2.5 \times 10^{-5}$ ) and low ( $2.5 \times 10^{-8}$ ). Ten replicates of test data Y were run to obtain Me. In the test data Z, with 100 Ne and 100 historical generations with mutation rate of $2.5 \times 10^{-8}$ , in the last generation, the population size was increased to 3000 and out of 3000 genotyped individuals, 2000 were included in the reference and 1000 in validation population. Accuracy was obtained as Pearson's correlation between TBV and GEBV. 35 replications were carried out for test data Z. Thirty chromosomes, each with 1 Morgan long, were used.

**Table S2:** Validation for the RR design simulated data with the theory (Lee et al. 2017)

| Data simulated for main study of the manuscript (RR Design) |  |  |  |  |  |
| --- | --- | --- | --- | --- | --- |
| heritability | GBLUP | G500 | G1000 | G2000 | G9550 |
| 0.10 | Expected accuracy with Ne=358 | 0.234 | 0.322 | 0.434 | 0.725 |
|  | Observed accuracy with GBLUP | 0.22±0.02 | 0.34±0.01 | 0.44±0.01 | 0.64±0.01 |
| 0.30 | Expected accuracy with Ne=358 | 0.385 | 0.508 | 0.641 | 0.877 |
|  | Observed accuracy with GBLUP | 0.38±0.01 | 0.49±0.01 | 0.61±0.01 | 0.79±0.01 |
| 0.50 | Expected accuracy with Ne=358 | 0.474 | 0.606 | 0.733 | 0.920 |
|  | Observed accuracy with GBLUP | 0.47±0.01 | 0.59±0.01 | 0.70±0.01 | 0.86±0.00 |

RR: No selection and random mating design; G500: GBLUP with 25% close male relatives (125) from 6<sup>th</sup> to 9<sup>th</sup> generations in reference; G1000: GBLUP with 50% close male relatives (250) from 6<sup>th</sup> to 9<sup>th</sup> generations in reference; G2000: GBLUP with 100% close male relatives (500) from 6<sup>th</sup> to 9<sup>th</sup> generations in reference; G9550: GBLUP with 9550 relatives from complete pedigree genotyped and used in reference. Observed estimates are provided with standard error. Thirty chromosomes, each with 1 Morgan long, were used.

**Table S3:** Number of Large QTL in validation population (Generation 10) explaining per cent variance

| Design<br>($h^2=0.30$) | QTL-model | 1% | 5% | 10% | 25% |
| --- | --- | --- | --- | --- | --- |
| **RR** | 90 | 18.08±0.70 | 5.12±0.35 | 1.96±0.16 | 0.32±0.10 |
|  | 990 | 18.6±0.60 | 0.36±0.11 | nil | nil |
|  | 60000 | Nil | nil | nil | nil |
| **SR** | 90 | 15.88±0.72 | 2.68±0.28 | 0.68±0.22 | 0.04±0.04 |
|  | 990 | 17.6±0.68 | 0.20±0.08 | 0.12±0.08 | nil |
|  | 60000 | Nil | nil | nil | nil |
| **SA** | 90 | 13.44±0.62 | 1.60±0.26 | 0.12±0.09 | nil |
|  | 990 | 13.76±2.75 | 0.16±0.07 | nil | nil |
|  | 60000 | Nil | nil | nil | nil |

**Table S4:**
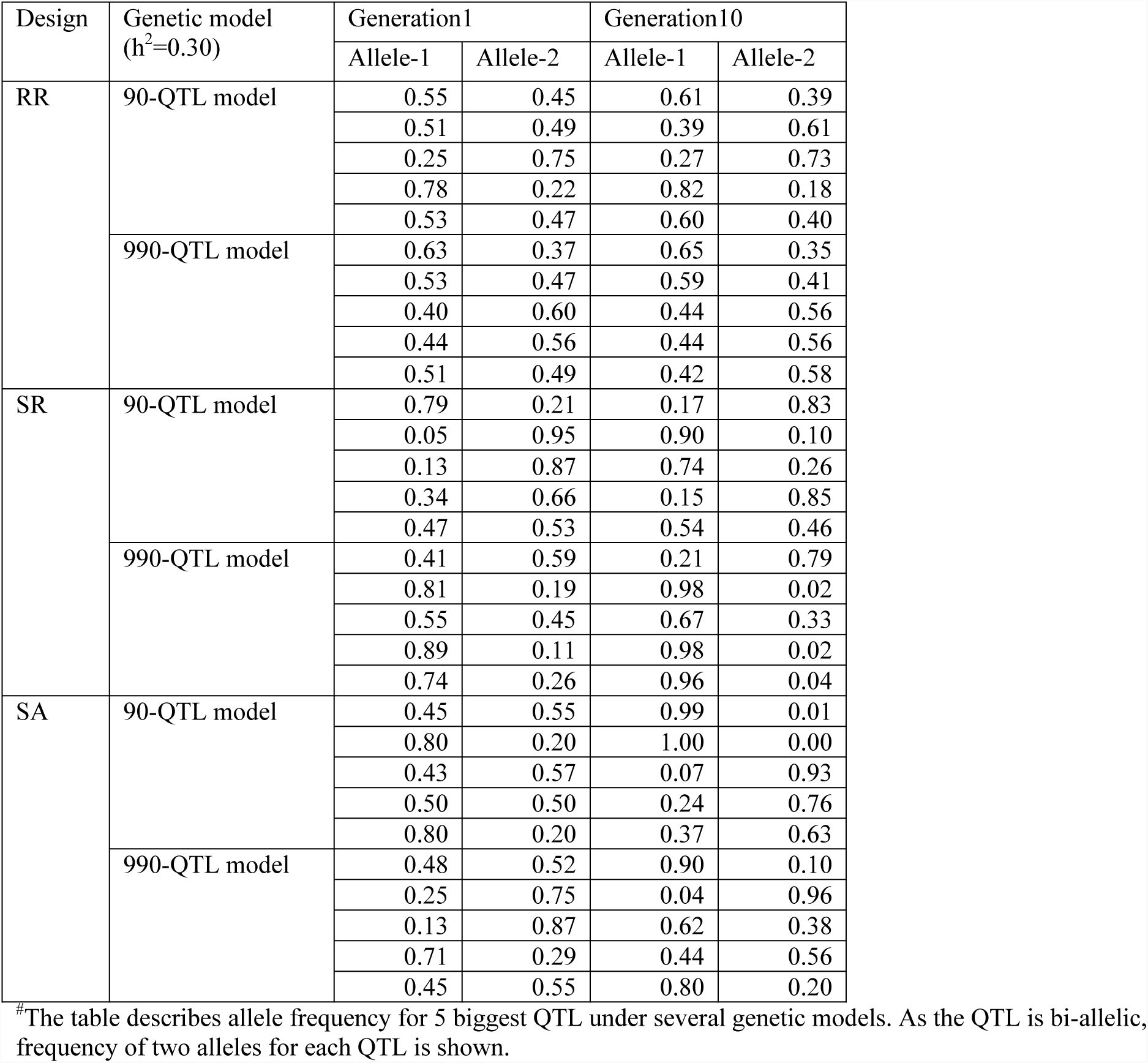
^#^Allele frequency changes for genetic model in 10 generations

| Design | Genetic model<br>($h^2=0.30$) | Generation1 | | Generation10 | |
| --- | --- | --- | --- | --- | --- |
|  |  | Allele-1 | Allele-2 | Allele-1 | Allele-2 |
| RR | 90-QTL model | 0.55 | 0.45 | 0.61 | 0.39 |
|  |  | 0.51 | 0.49 | 0.39 | 0.61 |
|  |  | 0.25 | 0.75 | 0.27 | 0.73 |
|  |  | 0.78 | 0.22 | 0.82 | 0.18 |
|  |  | 0.53 | 0.47 | 0.60 | 0.40 |
|  | 990-QTL model | 0.63 | 0.37 | 0.65 | 0.35 |
|  |  | 0.53 | 0.47 | 0.59 | 0.41 |
|  |  | 0.40 | 0.60 | 0.44 | 0.56 |
|  |  | 0.44 | 0.56 | 0.44 | 0.56 |
|  |  | 0.51 | 0.49 | 0.42 | 0.58 |
| SR | 90-QTL model | 0.79 | 0.21 | 0.17 | 0.83 |
|  |  | 0.05 | 0.95 | 0.90 | 0.10 |
|  |  | 0.13 | 0.87 | 0.74 | 0.26 |
|  |  | 0.34 | 0.66 | 0.15 | 0.85 |
|  |  | 0.47 | 0.53 | 0.54 | 0.46 |
|  | 990-QTL model | 0.41 | 0.59 | 0.21 | 0.79 |
|  |  | 0.81 | 0.19 | 0.98 | 0.02 |
|  |  | 0.55 | 0.45 | 0.67 | 0.33 |
|  |  | 0.89 | 0.11 | 0.98 | 0.02 |
|  |  | 0.74 | 0.26 | 0.96 | 0.04 |
| SA | 90-QTL model | 0.45 | 0.55 | 0.99 | 0.01 |
|  |  | 0.80 | 0.20 | 1.00 | 0.00 |
|  |  | 0.43 | 0.57 | 0.07 | 0.93 |
|  |  | 0.50 | 0.50 | 0.24 | 0.76 |
|  |  | 0.80 | 0.20 | 0.37 | 0.63 |
|  | 990-QTL model | 0.48 | 0.52 | 0.90 | 0.10 |
|  |  | 0.25 | 0.75 | 0.04 | 0.96 |
|  |  | 0.13 | 0.87 | 0.62 | 0.38 |
|  |  | 0.71 | 0.29 | 0.44 | 0.56 |
|  |  | 0.45 | 0.55 | 0.80 | 0.20 |

<sup>#</sup>The table describes allele frequency for 5 biggest QTL under several genetic models. As the QTL is bi-allelic, frequency of two alleles for each QTL is shown.

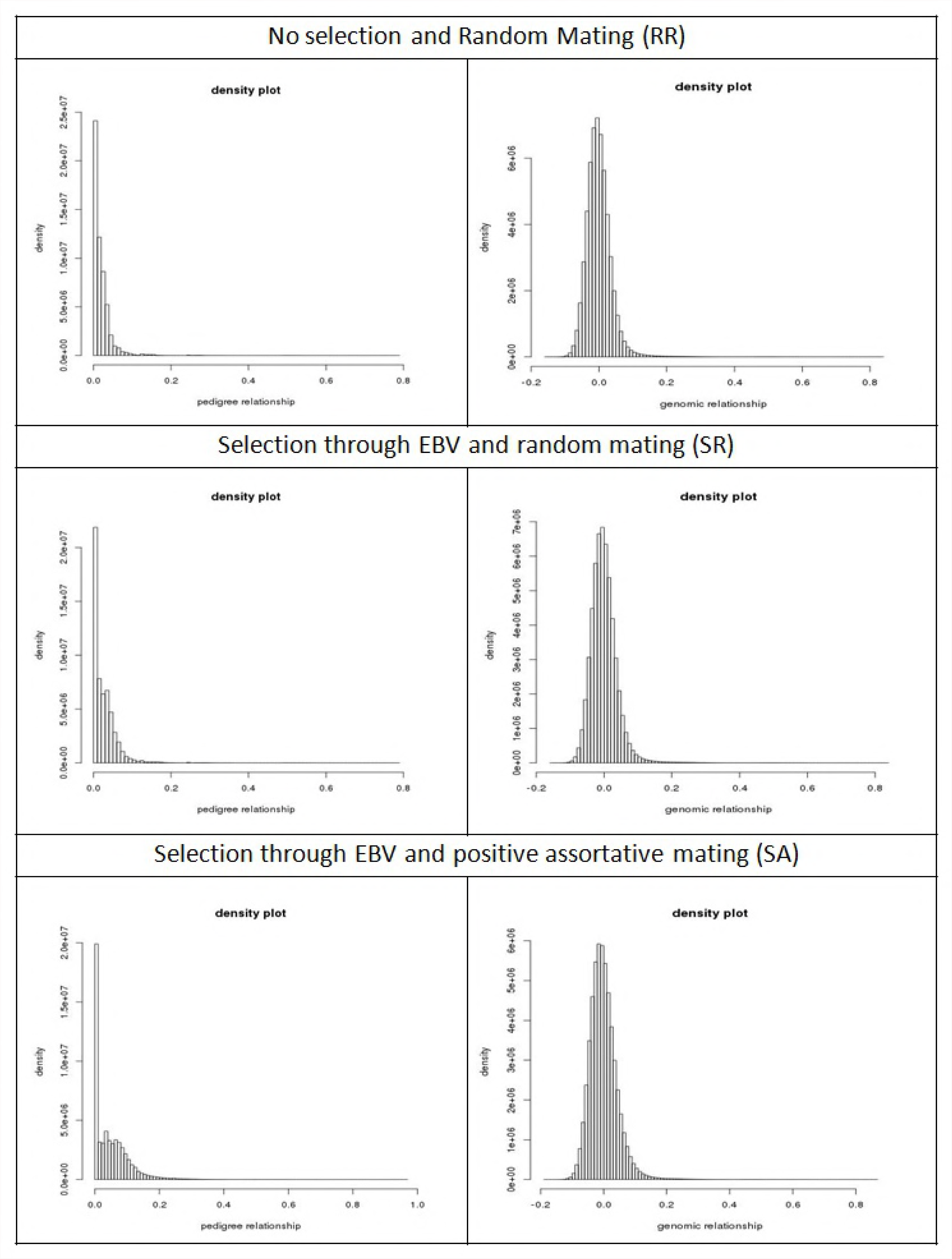

